# A pair-rule-like transcription network coordinates neural tube closure in a proto-vertebrate

**DOI:** 10.1101/2025.06.18.660479

**Authors:** Gabrielle Östlund-Sholars, Laurence A. Lemaire, Michael S. Levine

## Abstract

Neural tube closure (NTC) is a conserved morphogenetic process in chordates, during which the neural plate folds and fuses to form a closed neural tube. While the mechanical forces and signaling pathways governing NTC have been characterized in vertebrates, the transcriptional programs coordinating cell behaviors during closure remain less understood. Here, we identify a transcriptional circuit involving *Lmx1, Cdkn1b*, and *Msx* that regulates dorsal midline dynamics during NTC in the tunicate *Ciona*. High-resolution HCR *in situ* hybridization reveals that *Lmx1* is dynamically enriched at the zippering point and advances in a posterior-to-anterior transcription wave, while *Msx* is downregulated in the same region, marking a transition from early neural patterning to morphogenetic execution. As closure progresses, *Lmx1* and *Cdkn1b* exhibit complementary, alternating expression at the dorsal midline, resembling a pair-rule-like pattern. Misexpression studies show that *Lmx1* promotes proliferation and activates its own expression, whereas *Cdkn1b* limits proliferation and impedes closure. Single-cell RNA-seq reveals transcriptionally distinct dorsal neural populations enriched for *Lmx1* or *Cdkn1b*, supporting spatially organized cell-cycle states. These findings suggest that a transcriptional switch from *Msx* to *Lmx1*, followed by spatially alternating *Lmx1* and *Cdkn1b* activity, coordinates proliferation and neural fold fusion during NTC. This mechanism may represent a general strategy for regulating epithelial remodeling in animal embryos.

## Introduction

Neural tube closure (NTC) is a fundamental morphogenetic process in chordate development, during which the neural plate folds and fuses to form a hollow tube that gives rise to the central nervous system (Moon and Xiong, 2022; Nikolopoulou et al., 2017). In tunicates, this process proceeds through coordinated steps of neural plate bending, elevation of the neural folds, and a unidirectional posterior-to-anterior “zippering” of the neural folds across the dorsal midline (Hashimoto and Munro, 2019; Hashimoto et al., 2015), similar to the multiple closure points of vertebrates. This process relies on the coordinated integration of mechanical forces and signaling cues from both neural and surrounding tissues to ensure timely and ordered closure (Nikolopoulou et al., 2017). Disruptions in NTC can lead to neural tube defects (NTDs), which are among the most prevalent congenital malformations in humans (Kancherla, 2023; Avagliano et al., 2019; Greene and Copp, 2014), including conditions such as spina bifida and anencephaly.

Recent studies highlight the importance of spatiotemporal control of cell proliferation during NTC (Ogura et al., 2011). In vertebrates, spatial differences in the cell-cycle state influence epithelial stiffness, tension generation, and tissue fluidity, ultimately shaping morphogenetic outcomes (Bocanegra-Moreno et al., 2023). For instance, decreased proliferation can impair convergent extension, while excessive proliferation can cause overcrowding and mechanical jamming (Bocanegra-Moreno et al., 2023; Kim et al., 2021). However, how these proliferative dynamics interface with patterning and cell fate specification remains difficult to resolve in vertebrates, in part because changes in proliferation inherently alter total cell number. The ascidian *Ciona* provides a simple system for studying this process due to its invariant cell lineages and small cell numbers.

There are striking parallels between NTC in *Ciona* and vertebrates (Sasakura et al., 2012; Dahlberg et al., 2009). During neurulation, mitotic divisions propagate in a posterior-to-anterior wave across the epidermis, synchronized with neural fold fusion. This mitotic wave is transcriptionally patterned via *GATAb* and *AP-2* acting on the cell-cycle regulator *cdc25* (Ogura and Sasakura, 2016). This mechanism establishes cell-cycle compensation in the epidermis to ensure seamless fusion at the neural/epidermal (Ne/Epi) boundary. Cell-cycle compensation refers to the local acceleration or delay of cell divisions to equalize cell numbers across regions undergoing morphogenetic strain. Dysregulation of compensation prevents the progression of neural fold fusion (Ogura et al., 2011). These findings highlight how transcriptional regulation of cell-cycle dynamics facilitates coordinated epithelial remodeling (Ogura and Sasakura, 2016). In this context, cell-cycle inhibitors such as *Cdkn1b* (cyclin-dependent kinase inhibitor; p27^Kip1) emerge as important modulators of local proliferation (Treen et al., 2023b). While compensatory cell-cycle regulation has been studied in the overlying epidermis, it has not been characterized in the underlying neural cells undergoing neural tube zippering (Ogura and Sasakura, 2016).

In *Ciona*, zippering is driven by patterned actomyosin contractility and junctional exchange in dorsal neural tube cells. This behavior depends on local cytoskeletal asymmetry and cell intercalation (Hashimoto and Munro, 2019; Hashimoto et al., 2015). These findings link mechanical force generation to regional cell behaviors, raising the possibility that transcriptional patterning directly coordinates morphogenetic behaviors during neurulation. Transcription factors that simultaneously influence proliferation and cell fate likely contribute to coordinating proliferation and cell behaviors during NTC (Kim et al., 2022; Harris and Juriloff, 2010, 2007). LIM-homeodomain transcription factors, including LMX1A and LMX1B, are expressed in the dorsal midline of the vertebrate neural tube and are essential for roof plate specification and therefore dorsoventral patterning (Mishima et al., 2009; Chizhikov and Millen, 2004). These genes regulate the expression of morphogens such as *Wnt* and *BMP* and are necessary for the organization of dorsal midline cells and proper neural tube morphology (Lehr et al., 2025; Yan et al., 2011; Mishima et al., 2009; Riddle et al., 1995). In mice, spontaneous mutations in *Lmx1a* result in the *Dreher* phenotype, characterized by roof plate loss and cerebellar hypoplasia, underscoring its role in proper dorsal central nervous system patterning (Millonig et al., 2000). In both vertebrates and *Ciona, Msx* (muscle segment homeobox) genes mark the dorsal neural plate border and have been implicated in early neural patterning (Ramos and Robert, 2005; Imai et al., 2004), suggesting that multiple transcriptional programs contribute to dorsal midline organization.

Here, we employ single-cell methods to investigate the role of *Lmx1* in NTC in *Ciona*. We show that *Lmx1* is dynamically expressed in a posterior-to-anterior transcription wave at the advancing edge of closure, where it promotes cell proliferation and positively autoregulates its own transcription. Notably, we find that *Lmx1* might also activate its own expression in neighboring cells, raising the possibility of non-cell-autonomous expansion of the proliferative domain. However, we cannot exclude the possibility that prior *Msx* activity in some of these cells may have primed them for subsequent *Lmx1* activation. *Lmx1* and *Msx* are co-expressed in neural cells at the dorsal midline in early neurulation. As closure progresses, *Lmx1* becomes enriched at the zippering point while *Msx* is selectively downregulated, producing a locally exclusive pattern. This transition marks a shift from *Msx* -driven neural patterning to *Lmx1* -mediated morphogenesis. As closure advances, *Lmx1* and *Cdkn1b* adopt a complementary, alternating expression pattern along the neural folds. Functional perturbations reveal that both excessive and insufficient proliferation of future midline cells disrupt closure, underscoring the importance of spatially patterned proliferation. High-resolution imaging and single-cell RNA-seq confirm transcriptionally distinct dorsal neural populations. Together, these findings reveal a pair-rule-like transcriptional logic coordinating proliferation and morphogenesis, suggesting a novel mechanism by which alternating transcriptional states regulate epithelial remodeling. Our work identifies a previously uncharacterized role for *Lmx1* in morphogenesis, positioning it as a key node linking patterning, proliferation, and cell behavior in a chordate model of NTC.

## Results

### *Lmx1* expression advances in a posterior-to-anterior transcription wave during NTC

The *Ciona* genome contains two copies of *Lmx1* (*Lmx1* and *Lmx1-related*), both orthologous to the human paralog LMX1B (José-Edwards et al., 2011). Similar to their vertebrate counterparts, both *Ciona Lmx1* genes encode proteins with two tandem N-terminal LIM domains (each with a double zinc finger motif) and a C-terminal DNA-binding homeodomain (German et al., 1992). While *Lmx1-related* is expressed ventrally in the developing notochord (Negrón-Piñeiro et al., 2024), *Lmx1* is expressed dorsally in the cells of the roof plate that zipper during NTC (Cao et al., 2019; Imai et al., 2004). In vertebrates, *Lmx1a/b* are essential for dorsal midline patterning and roof plate formation (Millonig et al., 2000; Riddle et al., 1995).

Previous studies identified *Lmx1* expression along the future dorsal midline during neurulation, but with limited spatial and temporal resolution (Ishida and Satou, 2024; Imai et al., 2006, 2004). Using high-resolution HCR *in situ* hybridizations of precisely staged *Ciona* embryos, we found that *Lmx1* transcripts are not uniformly distributed but instead exhibit a posterior-to-anterior transcription wave during NTC (Fig. 1A-H). *Lmx1* is first detectable at the early gastrula stage at the neural plate border (*Fig. S1*), peaks toward the end of neurulation (Fig. 1M), and subsequently declines. A ridgeline plot of the normalized x- and y-coordinates of *Lmx1* -expressing nuclei along the anteroposterior axis reveals a forward-shifting wave of nuclear transcripts (Fig. 1L). Notably, *Lmx1* expression is the highest at the zippering point and in anterior neural cells poised to fuse but reduced in posterior cells that have already undergone closure (Fig. 1N-Q). This transcription wave coincides with the progression of zippering, suggesting that *Lmx1* activation either precedes or accompanies neural fold fusion and is tightly coupled to the morphogenetic events of zippering.

**Figure 1.**
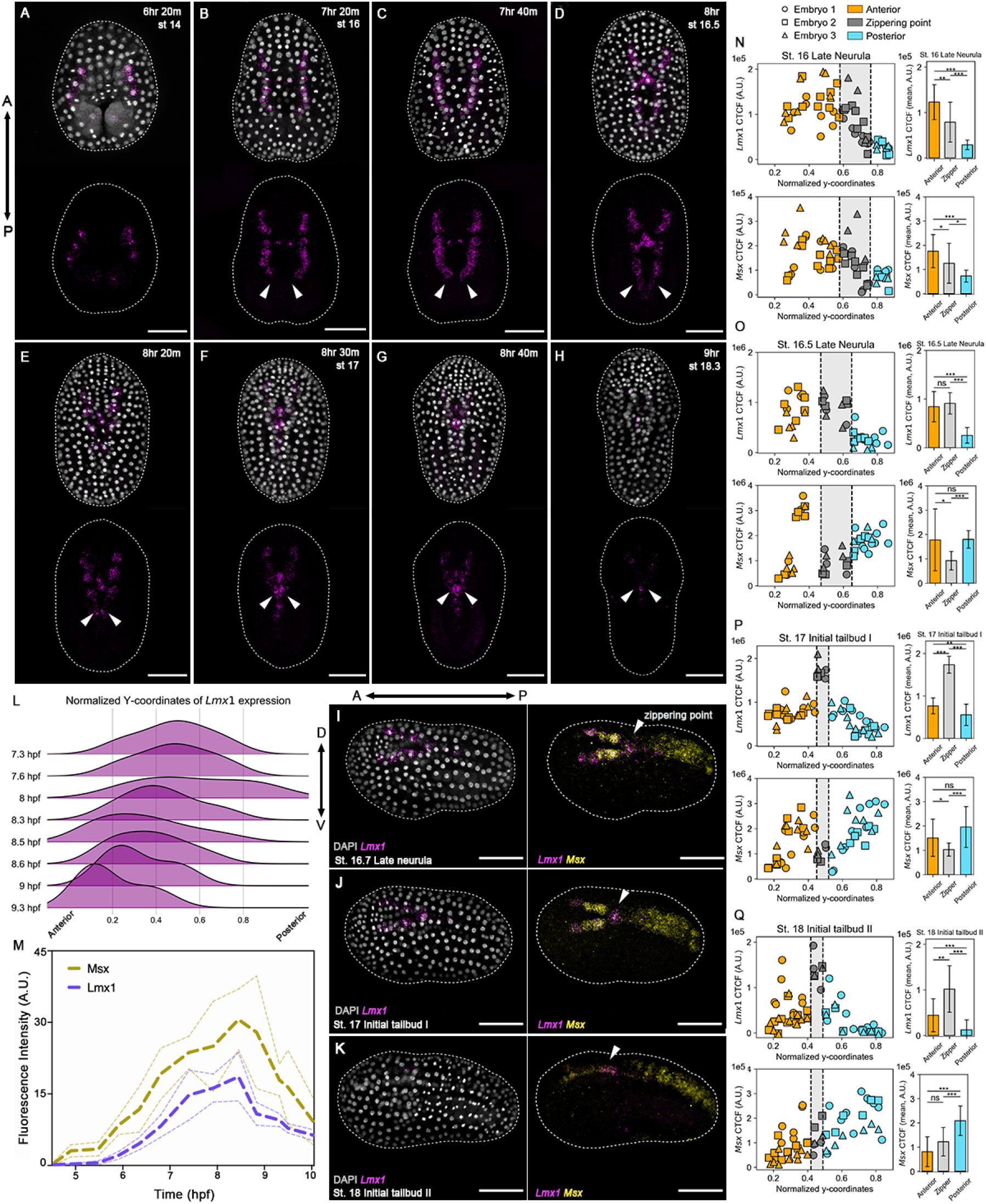
* *Lmx1* exhibits a posterior-anterior transcription wave during neural tube zippering. (A–H) Representative embryos showing HCR *in situ* hybridization signals for the *Lmx1* probe (magenta); (I–K) *Msx* probe (yellow); nuclei were stained with DAPI (gray); during neurulation. *Lmx1* is expressed in the intercalating cells at the head of the zipper and in the cells anterior to the zipper. (I–K) demonstrate the inverse expression pattern of *Lmx1* and *Msx* from 8–9 hpf. (L) Ridgeline plot showing the y-coordinates of *Lmx1* expression normalized to the embryo bounds for time points shown in (A–H), demonstrating the progression of *Lmx1* expression from posterior (right) to anterior (left) throughout neurulation (7.3–9.3 hpf). (M) Quantification of active transcription levels from *Lmx1* and *Msx* throughout gastrulation and neurulation (4.5–10 hpf); the solid line indicates the mean (n = 5 embryos per time point), and the scale bars indicate the standard deviation. (N–Q) Corrected total cell fluorescence (CTCF) for *Lmx1* and *Msx* HCR signals in dorsal midline cells during NTC, plotted relative to each cell’s y-coordinate normalized to the embryo bounds (n = 3 embryos per time point). Data were extracted from maximum intensity projections, with each point representing a single cell, and reveal that *Lmx1* is locally enriched at the zippering point (gray), whereas *Msx* is downregulated in the same region. (N–Q) bar graphs showing the mean CTCF for cells anterior (orange) of the zipper, at the zippering point (gray), and cells posterior (light blue) of the zipper. Error bars indicate the standard deviation for each group, *** p-value *<* 0.001, ** p-value *<* 0.01, * p-value *<* 0.05, n.s. ≥ 0.05. Developmental stages and hpf are indicated in the photographs. Brightness and contrast of images were adjusted linearly. Arrows (white) indicate the zippering point. Dashed lines (white) represent the embryo outline. Numbers of embryos examined: n = 15 per time point over n = 5 experiments. Scale bar: 50 μm.

We next examined the expression of *Msx*, a conserved dorsal neural plate border marker in both *Ciona* and vertebrates (Ramos and Robert, 2005; Imai et al., 2004). During early neurulation, *Msx* and *Lmx1* are broadly co-expressed along the dorsal midline (Fig. 1N; *Fig. S1A-G, Fig. S2A*). However, as the neural folds elevate and zipper, their expression becomes spatially refined (Fig. 1I-K). At the zippering point, *Lmx1* becomes enriched (*Fig. S2B-D*) while *Msx* is selectively downregulated (*Fig. S2B-C*), resulting in a locally exclusive pattern (Fig. 1O-Q, *Fig. S1H-L, Fig. S2B-C*). This transition indicates a spatially coordinated interplay between these transcriptional programs. The temporal handoff between *Lmx1* and *Msx* might reflect a regulatory switch, where we move from *Msx* -guided dorsal patterning to *Lmx1* - mediated morphogenesis.

### *Lmx1* promotes cell proliferation at the dorsal midline

To test the function of *Lmx1* during NTC, we overexpressed *Lmx1* under the control of the *Msx* enhancer. This led to an expanded dorsal midline domain (Fig. 2A-D), significantly increased numbers of *Lmx1* - (Fig. 2E) and *Msx* -reporter-positive nuclei (Fig. 2F), and impaired zippering (Fig. 2G-I). Phalloidin staining revealed disruptions to apical seam organization (Fig. 2J-L), suggesting that elevated cell density interferes with neural fold fusion, likely by perturbing the spatial constraints necessary for coordinated cell movements.

**Figure 2.**
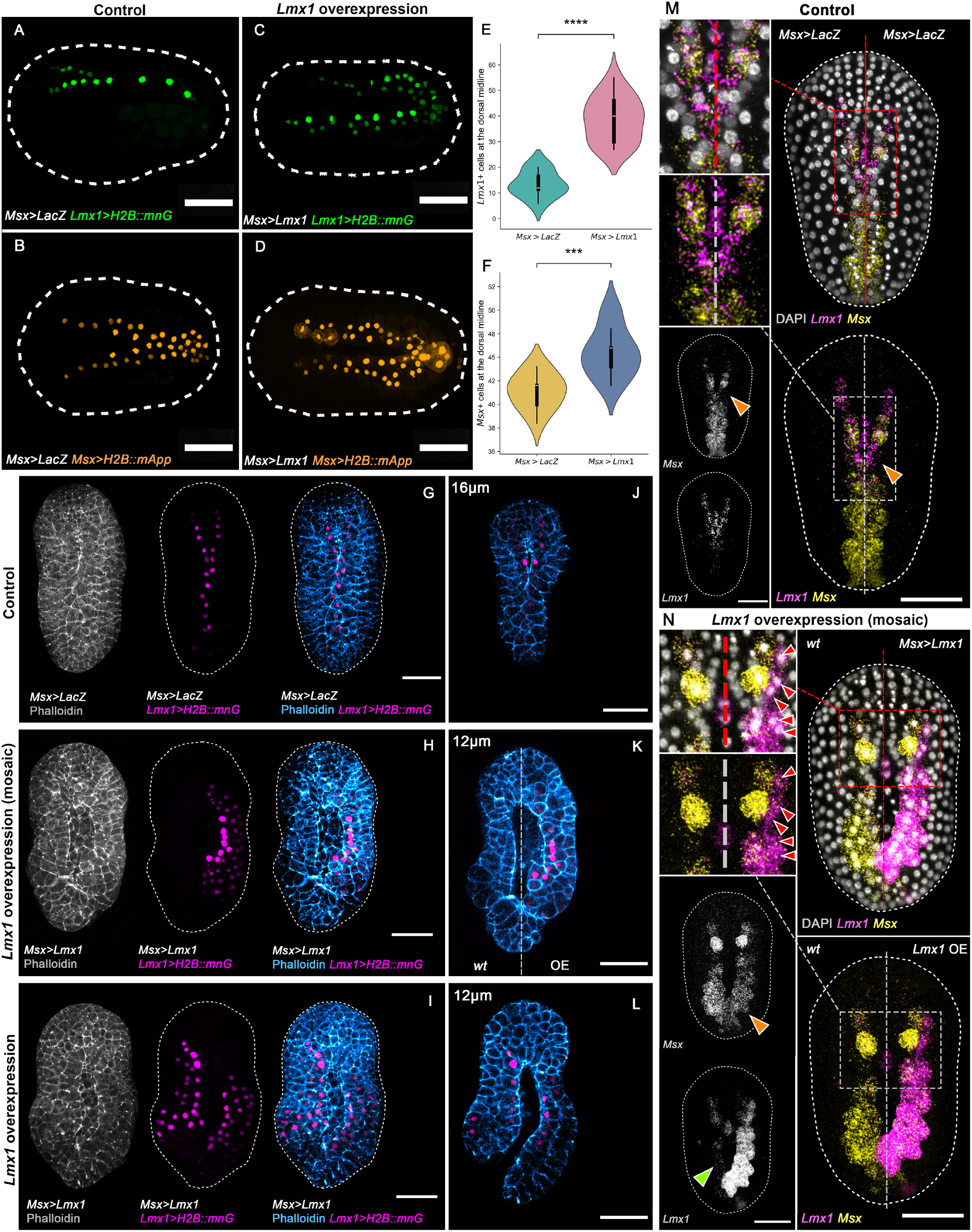
* *Lmx1* autoregulates and promotes cell proliferation during neural tube closure. (A-D) Representative initial tailbud I images of embryos expressing (A and B) *LacZ* under the regulatory sequence of *Msx* (*Msx>LacZ*), *Lmx1* fused to nuclear reporter *H2B::mnG* (*Lmx1>H2B::mnG*), green, (A), and *Msx* fused to nuclear reporter *H2B::mApple* (*Msx>H2B::mApp*), orange, (B). (C and D) Representative initial tailbud I embryos *Lmx1* under the regulatory sequence of *Msx* (*Msx>Lmx1*), *Lmx1* fused to nuclear *mnG* (*Lmx1>H2B::mnG*), green, (C), and *Msx* fused to nuclear *mApp* (*Msx>H2B::mApp*), orange, (D). (E-F) Violin plots with box plots of the number of dorsal midline cells expressing the (E) *Lmx1* nuclear reporter (*Lmx1>H2B::mnG*) or (F) the *Msx* nuclear reporter (*Msx>H2B::mApp*) at the initial tailbud I stage. (t-test, independent samples). (E) **** p-value = 4.1 × 10^*−*7^, (F) *** p-value = 3.7 × 10^*−*4^. (G-I) Representative initial tailbud I images of embryos expressing (G,J) *LacZ* under the regulatory sequence of *Msx* (*Msx>LacZ*), *Lmx1* fused to nuclear reporter *H2B::mnG* (*Lmx1>H2B::mnG*), magenta, and F-actin, gray and blue. (H-I, K-L) Representative initial tailbud I embryos expressing *Lmx1* under the regulatory sequence of *Msx* (*Msx>Lmx1*), (H,K) mosaic expression of *Msx>Lmx1* and *Lmx1>H2B::mnG* is restricted to the right side of the embryo. Photographs are maximum intensity projections of Z-projected image stacks overlaid in pseudocolor. (J-K) Representative images of optical slices at depths (D) 16 μm, and (E-F) 12 μm from the top of the embryo (dorsal side), from Z-projected image stacks in images (G-I), highlighting the effects on NTC. Numbers of embryos examined: n = 20 per experiment (n = 3). (M-N) Representative initial tailbud I images of embryos showing HCR *in situ* hybridization signals for the *Lmx1* probe (magenta) and *Msx* probe (yellow). Nuclei were stained with DAPI (gray). (M) Representative image of embryo expressing *LacZ* under the regulatory sequence of *Msx* (*Msx>LacZ*), and (N) mosaic overexpression of *Lmx1* under the regulatory sequence of Msx (*Msx>Lmx1*) on the right side of the embryo. Arrows indicate (1) *Msx* repression (orange) coinciding with *Lmx1* enrichment, (2) loss of the *Lmx1* transcription wave (green) coinciding with loss of zippering, and ectopic expression of *Lmx1* under the control of the *Msx* regulatory sequence (*Msx>Lmx1*) leads to activation of *Lmx1* in neighboring cells (red) outside of the *Msx* expression domain at the time of analysis, consistent with its ability to act non-cell-autonomously. However, we cannot rule out the possibility that these cells may have transiently expressed *Msx* earlier, contributing to their responsiveness *Lmx1* autofeedback. Brightness and contrast of images were adjusted linearly. Dashed lines (white) represent the embryo outline. Numbers of embryos examined: n = 20 per experiment (n = 3). Scale bar: 50 μm.

Ectopic *Lmx1* expression led to activation of an *Lmx1* reporter gene in the dorsal epidermis, outside of its normal domain of action (Fig. 2G-L). This indicates that *Lmx1* is sufficient to activate its own enhancer in a cell-autonomous manner and thus drive autoregulation. This autoregulatory capacity may reinforce *Lmx1* expression in zippering cells and sustain the proliferative state. Importantly, in mosaic embryos, we observed that *Msx>Lmx1* overexpression led to activation of *Lmx1* in neighboring cells that did not express *Msx* at the time of analysis (Fig. 2N; red arrows), suggesting non-autonomous expansion of *Lmx1* expression. These findings support a model in which *Lmx1* not only reinforces its own expression through direct positive feedback but also amplifies its spatial domain through local cell-cell interactions. However, it is conceivable that transient, earlier *Msx* expression in these cells could also contribute to their capacity for *Lmx1* induction. This autoregulatory and spatially expansive behavior suggests that *Lmx1* sits at the core of a dynamic transcriptional circuit that integrates proliferation, patterning, and mechanical coordination during NTC.

In these same mosaic embryos, *Lmx1* overexpression led to local downregulation of *Msx* in *Lmx1* - overexpressing cells (Fig. 2M-N; orange arrows), consistent with repressive feedback from downstream components described in prior studies (Roure and Darras, 2016), and may suggest that *Lmx1* may participate directly or indirectly in repressing *Msx* expression to promote morphogenetic progression. Notably, the endogenous *Lmx1* transcription wave was disrupted (Fig. 2N; green arrow), suggesting that either the transcription wave is necessary for zippering or that ongoing zippering is required to maintain *Lmx1* activation dynamics. Taken together, these findings show that *Lmx1* reinforces its own expression cell-autonomously, and might also contribute to the expansion of the proliferative domain in a non-autonomous fashion. This behavior is consistent with a positive feedback loop operating within a bistable or excitable system (Zhao et al., 2023) and underscores the importance of precisely patterned *Lmx1* expression for robust morphogenesis.

### *Cdkn1b* restricts proliferation and contributes to proper zippering dynamics

Given that *Lmx1* might promote proliferation, we next asked whether *Cdkn1b* (p27^Kip1), a known cell-cycle inhibitor (Treen et al., 2023b), acts as a counterbalance during NTC. High-resolution HCR *in situ* hybridization enabled us to visualize the spatiotemporal dynamics of *Cdkn1b* expression throughout neurulation (*Fig. S3; Fig. S4*). We found that *Cdkn1b* is expressed globally throughout neurulation, with enrichment along the dorsal midline during anterior closure (*Fig. S4E-K*). Notably, analysis of individual confocal z-planes through the roof plate revealed a striking alternating pattern of *Cdkn1b* expression in neural cells at the dorsal midline, with adjacent cells showing either high or undetectable levels. This pattern mirrored that of *Lmx1* and suggested a spatially coordinated transcriptional program.

To test *Cdkn1b* function, we overexpressed it using the regulatory sequence of *Lmx1*, restricting expression to dorsal midline neural cells. Compared to *LacZ* controls, *Cdkn1b*-overexpressing embryos were shorter (Fig. 3A-D), had a reduced number of neural tube cells (Fig. 3E-F), and exhibited stalled NTC (Fig. 3G-H). Unlike *Lmx1* overexpression, which disrupted apical F-actin, *Cdkn1b*-overexpressing embryos retained apical seam integrity at the zippering point but showed reduced dorsal tissue (Fig. 3I-J). These observations imply that impaired closure in *Cdkn1b*-overexpressing embryos reflects insufficient cell number rather than defects in cytoskeletal organization. These findings reveal that neural fold fusion is constrained by a biophysical threshold of local cell density below or above which fusion fails to proceed efficiently and that *Cdkn1b* may help modulate this threshold. Together, these results point to a finely tuned, transcriptionally regulated coordination of cell-cycle dynamics in both neural and epidermal tissues during neurulation.

**Figure 3.**
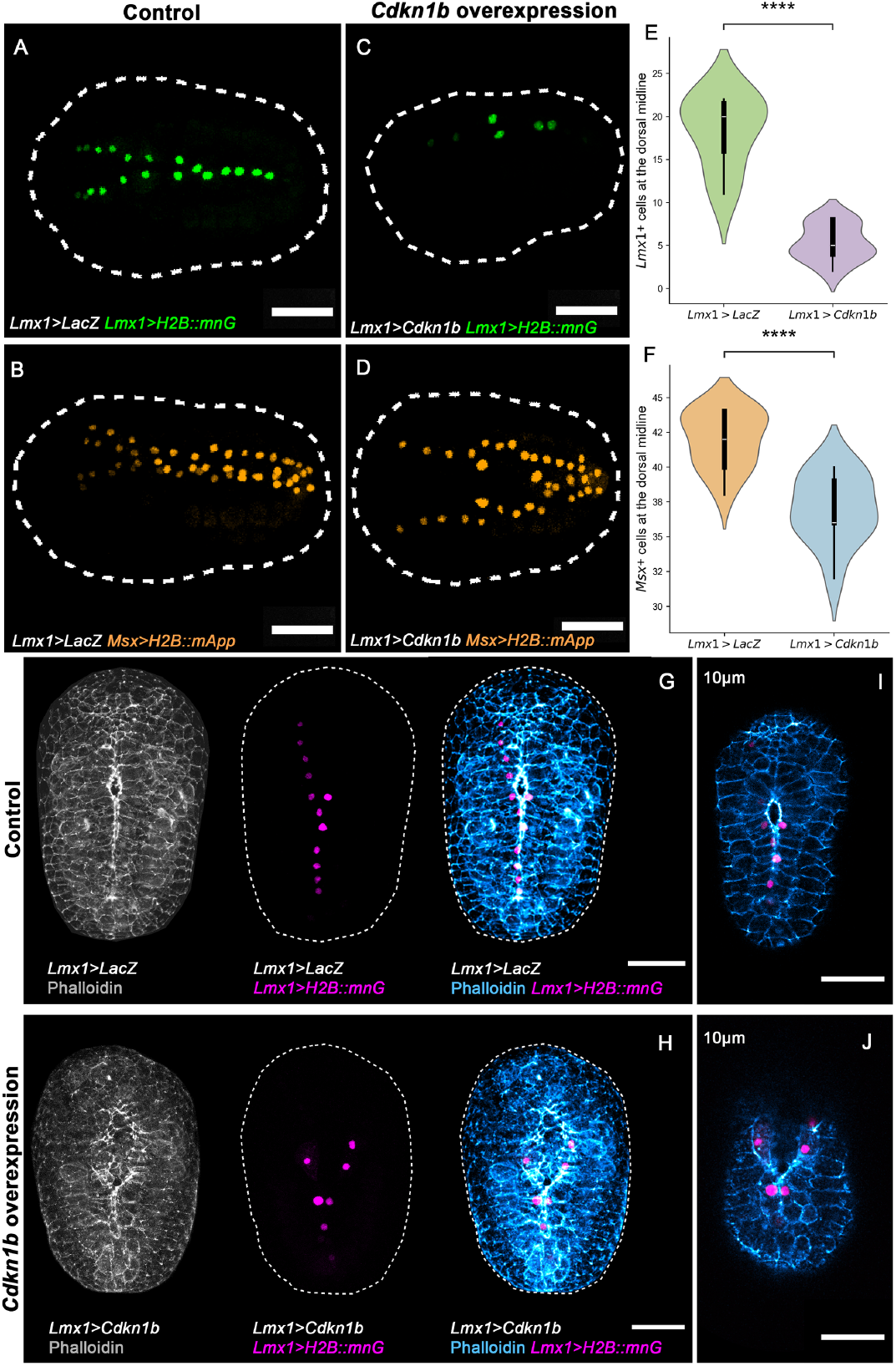
* *Cdkn1b* involvement in cell cycle regulation during neural tube closure. (A-D) Representative initial tailbud I images of embryos expressing (A and B) *LacZ* under the regulatory sequence of *Lmx1* (*Lmx1>LacZ*), *Lmx1* fused to nuclear reporter *H2B::mnG* (*Lmx1>H2B::mnG*), green, (A), and *Msx* fused to nuclear reporter *H2B::mApple* (*Msx>H2B::mApp*), orange, (B). (C and D) Representative initial tailbud I embryos expressing *Cdkn1b* under the regulatory sequence of *Lmx1* (*Lmx1>Cdkn1b*), *Lmx1* fused to nuclear *mnG* (*Lmx1>H2B::mnG*), green, (C), and *Msx* fused to nuclear *mApp* (*Msx>H2B::mApp*), orange, (D). (E-F) Violin plots with box plots of the number of dorsal midline cells expressing the (E) *Lmx1* nuclear reporter (*Lmx1>H2B::mnG*) or (F) the *Msx* nuclear reporter (*Msx>H2B::mApp*) at the initial tailbud I stage. (t-test, independent samples). (E) **** p-value = 9.3 × 10^*−*10^, (F) **** p-value = 1.7 × 10^*−*5^. Photographs are Z-projected image stacks overlaid in pseudocolor. (G-J) Representative initial tailbud I images of embryos expressing (G,I) *LacZ* under the regulatory sequence of *Lmx1* (*Lmx1>LacZ*), *Lmx1* fused to nuclear reporter *H2B::mnG* (*Lmx1>H2B::mnG*), magenta, and F-actin, gray and blue. (H, J) Representative initial tailbud I embryos expressing *Cdkn1b* under the regulatory sequence of *Lmx1* (*Lmx1>Cdkn1b*). Photographs are maximum intensity projections of Z-projected image stacks overlaid in pseudocolor. (I-J) Representative images of optical slices at a depth of 10 μm from the top of the embryo (dorsal side), from Z-projected image stacks in images (G-H), highlighting the effects on NTC. Brightness and contrast of images were adjusted linearly. Dashed lines (white) represent the embryo outline. Numbers of embryos examined: n = 20 per experiment (n = 3). Scale bar: 50 μm.

### *Lmx1* and *Cdkn1b* exhibit a pair-rule-like expression pattern during anterior NTC

To investigate the spatial relationship between *Lmx1* and *Cdkn1b* expression, we performed multiplex HCR *in situ* hybridizations. We found that these genes exhibit strikingly complementary expression patterns across the dorsal midline, with *Lmx1* and *Cdkn1b* localized to adjacent, non-overlapping cells (Fig. 4A-C). This alternating, pairrule-like motif was especially prominent in the anterior neural folds during active zippering (Fig. 4A-F), suggesting that *Lmx1* and *Cdkn1b* define transcriptionally distinct anti-proliferative (*Cdkn1b*) and permissive (*Lmx1*) regulatory regimes within a shared morphogenetic field. The emergence of this pattern coincides temporally with peak zippering behavior, raising the possibility that alternating transcriptional states serve to modulate cell density and behavior at the cellular scale, reminiscent of pair-rule patterning logic in other systems (Clark et al., 2019; Nüsslein-Volhard and Wieschaus, 1980).

**Figure 4.**
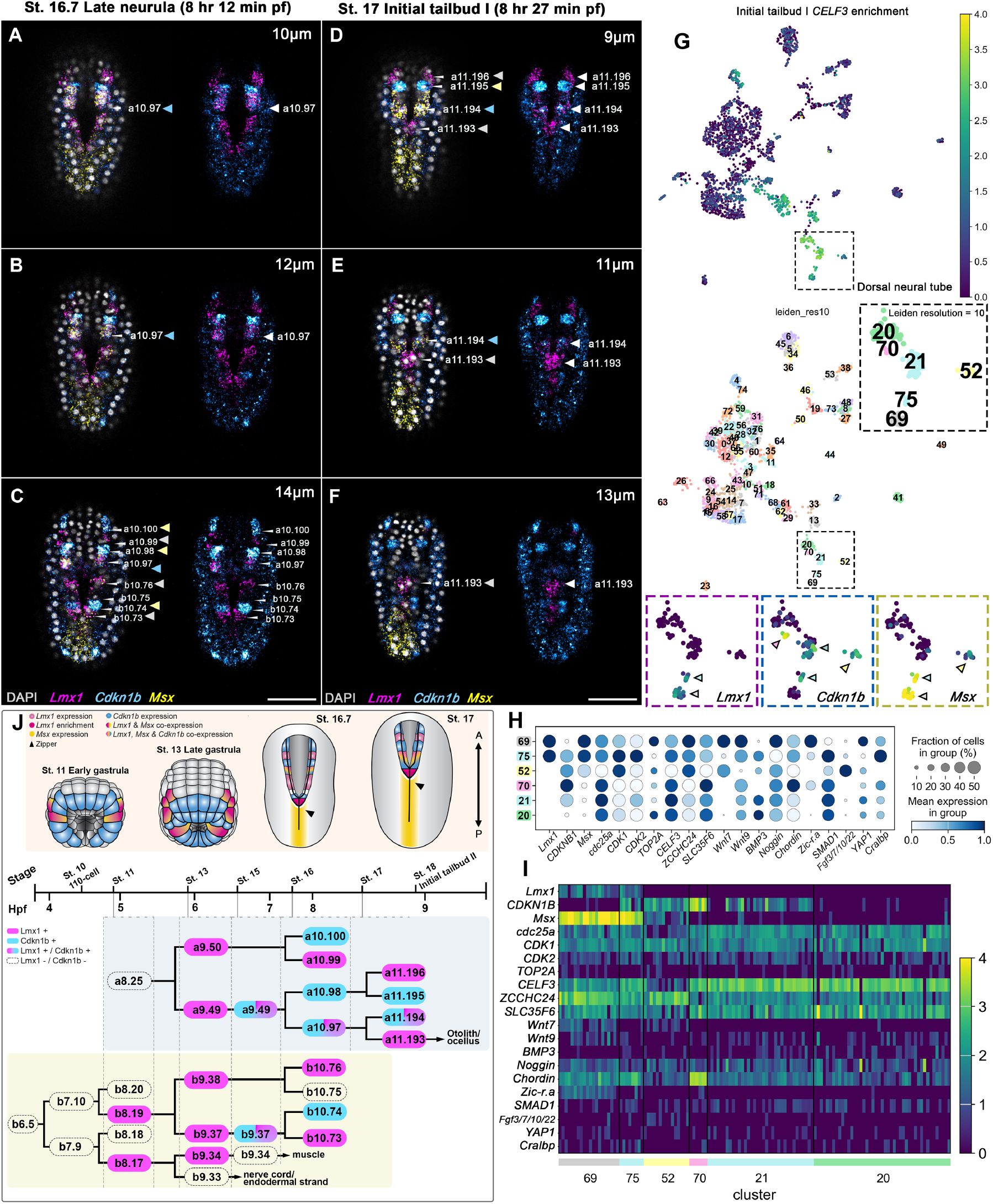
* *Lmx1* and *Cdkn1b* exhibit a pair-rule-like alternating expression during anterior neural fold fusion, suggesting patterned transcriptional control of neural fold fusion during closure. (A-C) Representative images of optical slices at depths (A) 10 μm, (B) 12 μm, and (C) 14 μm from the top of the embryo (dorsal side), from a Z-projected image stack of late neurula (8 hours 12 minutes pf) embryos that display the pair-rule-like expression pattern outlined in (J), where *Lmx1* and *Cdkn1b* show an alternating expression pattern in dorsal midline cells that are zippering during NTC. (D-F) Representative images of optical slices (D) 9 μm, (E) 11 μm, and (F) 12 μm from a Z-projected image stack of initial tailbud I (8 hours 27 minutes pf) embryos. Photographs are Z-projected image stacks overlaid in pseudocolor with HCR *in situ* hybridization signals for *Lmx1* probe (magenta), *Msx* probe (yellow), and *Cdkn1b* probe (cyan). Nuclei were stained with DAPI (gray). (G) Representative UMAP projection of single-cell transcriptomes of a representative initial tailbud I embryo colored by expression of selected dorsolateral neural marker gene *CELF3* (Copley et al., 2024) and Leiden clusters (resolution = 10). Expression levels are shown as a continuous color scale, while pastel hues represent cluster identities. The clusters representing the roof plate (20, 21, 52, 69, 70) are indicated in the subpanel. Expression levels for *Lmx1, Cdkn1b*, and *Msx* in respective clusters are shown in subpanels, indicating the transcriptionally defined cell populations corresponding to the pair-rule-like pattern observed *in vivo*. (H) Dot plot showing relative expression levels of selected genes across neural clusters shown in (G). Genes were chosen based on their relevance to neural identity, cellcycle regulation, and epithelial remodeling. (I) Heatmap of the same genes highlighted (H), illustrating expression patterns across individual cells within the neural lineage. Arrows (A-C) indicate the cells highlighted in magenta/cyan in the cell lineage (J) of the proximal (b8.17) and distal (b8.19) neural plate border cells, as well as in the cell lineage (a9.49) that gives rise to the rudimentary neural crest lineage. Brightness and contrast of images were adjusted linearly. Numbers of embryos examined: n = 15 per experiment (n = 5). Scale bars: 50 μm.

To test this idea, we analyzed previously published scRNA-seq data (Cao et al., 2019) from initial tailbud I embryos (Fig. 4G). Within the neural lineage, *Lmx1* and *Cdkn1b* expression mirrored the mutually exclusive patterns observed *in vivo*. High-resolution clustering (Leiden resolution = 10) further revealed three transcriptionally distinct dorsal neural populations (Fig. 4G). These include (1) *Lmx1+*/*Msx* + cells enriched for proliferative markers such as *cdc25a* and *CDK1* (Fig. 4H-I; cluster 69), (2) *Msx+*/*Cdkn1b+* cells enriched for differentiation and signaling markers such as *Fgf3/7/10/22* (Fig. 4H-I; cluster 52), and (3) a mixed population co-expressing all three (Fig. 4H-I; cluster 75) in addition to the ependymal marker *Cralbp*, which may indicate a transitional progenitor pool, potentially contributing to the rudimentary neural crest. This suggests a regulatory switch where cell-cycle exit and differentiation are tightly coupled to lineage identity.

*In vivo*, co-expression of *Lmx1, Msx*, and *Cdkn1b* was restricted to the a10.97 lineage (Fig. 4A-C, Fig. 4J), previously shown to give rise to the rudimentary neural crest lineage in *Ciona* (Todorov et al., 2024). Outside this lineage, *Lmx1* and *Cdkn1b* expression remained largely mutually exclusive, reinforcing the idea that their alternation reflects a functional division between proliferative and differentiating states. This complementary patterning may help buffer against mechanical or proliferative noise by distributing cell states in a spatially interdigitated fashion, thereby enhancing the robustness of zippering. Together, these findings reveal that a pair-rule-like transcriptional logic underlies dorsal midline organization during NTC.

## Discussion

This study identified a novel transcriptional framework that coordinates cell-cycle state and morphogenetic behavior during NTC in *Ciona*. We show that *Lmx1* functions as a proliferative transcription factor expressed in a dynamic posterior-to-anterior transcription wave at the zippering point. *Lmx1* positively autoregulates its own expression, expands the local proliferative domain, and is essential for proper neural fold fusion. In contrast, the cell-cycle inhibitor *Cdkn1b* acts as a spatially restricted brake on proliferation, expressed in an alternating pattern with *Lmx1*. This complementary expression balances cell density across the neural folds and the roof plate during anterior NTC.

A key feature of this system is a temporal handoff between *Msx* and *Lmx1* during NTC. Early in neurulation, *Msx* and *Lmx1* are co-expressed in the neural fold, but as closure progresses, *Msx* becomes selectively downregulated at the zippering point while *Lmx1* is upregulated. Our data indicate that this transition reflects not only passive downregulation of *Msx*, but also active repression. Our findings show that *Lmx1* overexpression leads to local downregulation of *Msx* (Fig. 2N; orange arrows), which is consistent with cell-autonomous negative feedback. This observation is aligned with previous gene regulatory network reconstructions in *Ciona* that identified widespread negative feedback onto *Msx* from its downstream targets (Roure and Darras, 2016). The emergence of *Lmx1* in *Msx* -negative cells marks a transition from early patterning to morphogenetic activity, where cells begin to proliferate and intercalate. This mutually exclusive expression indicates a transcriptional switch that gates entry into a proliferative, mechanically permissive state, positioning *Msx* -to-*Lmx1* transition as a key regulatory node in coordinating spatial organization during NTC.

The interleaved expression of *Lmx1* and *Cdkn1b* evokes the pair-rule patterning logic classically associated with arthropod segmentation (Zallen and Wieschaus, 2004; Irvine and Wieschaus, 1994). By alternating cycling and arrested cells, this transcriptionally patterned landscape may prevent mechanical jamming and allow more efficient fusion. The transient block on the cell-cycle due to the expression of *Cdkn1b* might be necessary to prevent cell division in front of the zippering point, which would lead to its arrest as observed with the adjacent epidermis (Ogura and Sasakura, 2016; Ogura et al., 2011). Computational models in other systems have previously shown that mechanical heterogeneity, typically achieved via differential tension and cell-cycle state, enhances robustness in tissue closure (Bocanegra-Moreno et al., 2023; Kim et al., 2021; Angelini et al., 2011). Our findings provide empirical evidence for such a mechanism in a chordate. It is possible that pair-rule segmentation in *Drosophila* and other arthropods establishes similar differential proliferative states to foster morphogenetic processes such as germband elongation (Clark et al., 2019).

Our perturbation data highlights the importance of spatially tuned proliferation. Overexpression of *Lmx1* leads to dorsal crowding, disrupted F-actin polarity, and failed zippering. This is consistent with mechanical jamming, a condition where excess cell density in conjunction with tension fluctuations inhibits coordinated movement due to overcrowding, reducing tissue fluidity (Sakamoto et al., 2023; Kim et al., 2021). Conversely, *Cdkn1b* overexpression reduces neural cell number and stalls NTC, despite preserved cytoskeletal organization. These embryos may experience mechanical under-loading, where insufficient traction or cell-cell contact prevents the generation of forces needed to drive zippering. This aligns with known previous observations of decreases in proliferation rate leading to a decline in cell rearrangements and tissue growth rate (Bocanegra-Moreno et al., 2023). These outcomes suggest that neural fold fusion efficiency operates within a “Goldilocks” zone of cell density, modulated by opposing cues.

Mechanistically, the *Lmx1-Cdkn1b* circuit might regulate both proliferation and biomechanical properties of zippering cells. The capacity of *Lmx1* to activate its own expression both cell-autonomously and potentially non-autonomously suggests a positive feedback loop stabilizing proliferative domains across the neural folds. However, we cannot rule out the possibility that prior transient *Msx* expression contributes to non-autonomous spatial expansion. Moreover, the mutual exclusivity of *Lmx1* and *Msx* at the zippering point further supports the idea of a bistable transcriptional switch coordinating morphogenetic transitions.

Single-cell RNA-seq analysis corroborates this framework, revealing three distinct dorsal neural transcriptional states linked to specialized roles during NTC. One cluster co-expressing *Lmx1* and *Msx* (Fig. 4H; cluster 69) also exhibits high expression levels of *Wnt7* /*9, Noggin, YAP1*, and cell-cycle drivers (*cdc25a, CDK1*, and *TOP2A*) but nearly undetectable levels of *Cdkn1b*, which is consistent with a transcriptional profile characteristic of proliferative and mechanosensitive fusing neural folds, forming one half of the pair-rule-like pattern where alternating cells drive closure. A second cluster, uniquely co-expressing *Lmx1, Cdkn1b*, and *Msx* (Fig. 4H; cluster 75) along with *Cralbp* and moderate levels of *Wnt9, Chordin*, and *SMAD1*, may represent a transitional progenitor pool. Its mixed expression of proliferative and inhibitory cell-cycle markers suggests bistability, potentially primed for fate transitioning rather than sustained division. The third *Cdkn1b* and *Msx* co-expressing cluster (Fig. 4H; cluster 52) is characterized by high *Fgf3/7/10/22* (the closest human gene is *FGF3*) expression and an absence of *Lmx1, Zic-r*.*a*, and *YAP1*, marking a transition from dorsal progenitor identity to a post-mitotic, differentiating neural fate. Although the expression of *CDK1* remains high, the reduced expression of *CDK2* and *TOP2A* suggests cells are withdrawing from the cell-cycle or entering arrest. This population likely represents the complementary half of the pair-rule architecture, contributing structural stability during NTC. Together, these data suggest a transcriptional landscape with intricate links between cell-cycle dynamics, patterning state, and differentiation.

### Summary

We report a striking pair-rule-like transcriptional logic co-ordinating proliferation and morphogenesis during NTC. The alternating expression of *Lmx1* and *Cdkn1b* delineates distinct proliferative regimes, while a temporal handoff from *Msx* to *Lmx1* marks the transition from early patterning to morphogenetic execution. This transcriptional architecture likely ensures robust neural fold fusion by spatially organizing mechanical forces and proliferation and may represent a conserved strategy across metazoans for integrating fate, behavior, and tissue mechanics.

## Methods

### *Ciona* handling, embryo collection, and electroporation

Adult *Ciona intestinalis* (type A; also called *Ciona robusta*) were obtained from M-Rep (San Diego, CA) and Marinus Scientific LLC (Long Beach, CA) and maintained in artificial seawater at 18 ^*°*^C. Eggs and sperm were harvested and dechorionated using a solution containing 60 mM NaOH, 1% (w/v) sodium thioglycolate, and 0.1% (w/v) Actinase E in artificial seawater for 10 minutes, as previously described (Christiaen et al., 2009b). Dechorionated eggs were then washed four times with artificial seawater before self-fertilization by incubation with sperm for 15 minutes, followed by two additional washes. All embryos used in this study are biological replicates.

One-cell stage embryos were transfected by electroporation with plasmid DNA, in accordance with a previously described method (Christiaen et al., 2009a). Transgenic embryos were cultured in artificial seawater on gelatincoated dishes at 18 ^*°*^C. until the desired developmental stage (Hotta et al., 2007). Embryos destined for cytoskeletal visualization were then fixed in MEM-PFA (4% paraformaldehyde, 100 mM MOPS pH 7.4, 500 mM NaCl, 1 mM EGTA, 2 mM MgSO_4_, and 0.05% Tween-20 in ultrapure H_2_O) for 30 minutes at room temperature, followed by three washes in PBST (phosphate buffer solution with 0.1% Tween-20). To visualize the cytoskeleton, embryos were incubated with Alexa Fluor 555-phalloidin (Abcam; 40X stock, used at 1:50 dilution) for 30 minutes and then washed three times with PBST.

### Molecular cloning and functional assays

The reporter *Lmx1>H2B::mnG* was previously described (Todorov et al., 2024). The *Msx* regulatory sequence published by Abitua and colleagues (Abitua et al., 2012) was subcloned into a *H2B::mApp* expression plasmid (Cao et al., 2019) using *NotI* and *AscI* restriction enzymes (New England Biolabs). Control constructs (*Lmx1>LacZ* and *Msx>LacZ*) were obtained by subcloning *Lmx1* and *Msx* regulatory sequences into *LacZ* expression plasmids (Fujiwara et al., 1998) using *NotI* and *AscI* restriction enzymes (New England Biolabs).

The regulatory sequences of *Lmx1* (2,029 bp; KH2012.C9:4348500-4350529) and *Msx* (2,446 bp; KH2012.C2:6131140-6133586) were PCR-amplified from the plasmids *Lmx1>H2B::mnG* and *Msx>H2B::mApp*. Reporter and overexpression constructs were generated via Gibson assembly using PCR-amplified fragments with designed overhangs (primer sequences provided in the *Supplementary Appendix*). To generate *Lmx1>GFP*, the *Lmx1* regulatory sequence was PCR-amplified from *Lmx1>H2B::mnG*, and the *GFP* coding sequence (717 bp) from the *TRE3G-GFP* vector (gift of Dr. Charles Ettensohn). For *Lmx1>Cdkn1b*, the *Lmx1* regulatory sequence was paired with the *Cdkn1b* coding sequence (915 bp) amplified from *Mesp>CkiB* (gift of Dr. Nicholas Treen). To create *Msx>Lmx1*, the *Msx* regulatory sequence was combined with the *Lmx1* coding sequence amplified from cDNA obtained by retrotranscription using Superscript II Reverse Transcriptase (Invitrogen) of 800 ng *Ciona* neurula RNA extracted using Qiagen RNA Extraction Kit. Transgenic constructs were assembled into circular plasmids using HiFi DNA Assembly (New England Biolabs) and introduced via electroporation at the one-cell stage.

All constructs were validated by sequencing, and all functional assays were performed at least twice with batches of embryos harvested from different animals. Gene identifiers (KY21 gene model; *Supplementary Appendix*) used in this study are as follows: KY21.Chr9.606 (*Lmx1*); KY21.Chr2.1031 (*Msx*); KY21.Chr2.18 (*Cdkn1b*).

### HCR *in situ* hybridization and imaging

Embryos were cultured in artificial seawater on gelatincoated dishes at 18 ^*°*^C until the desired developmental stage (Hotta et al., 2007) and then fixed in EGS fixative (1% formaldehyde, 100 mM HEPES, 500 mM NaCl, 2 mM MgSO_4_, 2 mM EGS [ethylene glycol bis(succinimidyl succinate)] in ultrapure H_2_O) and stored at -20 ^*°*^C prior to staining. Hybridization chain reaction (HCR) *in situ* hybridization was performed using reagents and protocols provided by Molecular Instruments (sea urchin protocol), with minor modifications: the probe-hybridization buffer contained 50% formamide, 5X Denhart’s, 0.01% (w/v) ss-DNA, 0.01% (w/v) yeast DNA, and 5X SSC (Treen et al., 2023a). The hairpin concentration was adjusted to 3 μM (2 μL hairpin in 100 μL amplification buffer), and hairpin amplification incubation time was shortened to 5 hours. DAPI (1 μg/mL, Thermo Fisher Scientific) was added post-hybridization to visualize nuclei. Embryos were washed in 5X SSCT and stored in PBST at 4 ^*°*^C prior to imaging. Probe sets (probe sequences in *Supplementary Appendix*) were designed by Molecular Instruments using the KY21 gene models: KY21.Chr9.606.v1.SL2-1 (*Lmx1*), KY21.Chr2.1031.v1.nonSL5-1 (*Msx*), and KY21.Chr2.18.v1.SL1-1 (*Cdkn1b*), compatible with amplifiers B1 (*Lmx1*), B3 (*Cdkn1b*), and B5 (*Msx*), respectively.

Samples were imaged using a Zeiss LSM 880 confocal microscope (Carl Zeiss) with a 20x objective, using Zen Black acquisition software. Z-stacks were acquired at 1 μm intervals, with a pixel size of 0.198 x 0.198 μm, and fluorophores were excited using 405, 488, 514, 561, and 633 nm lasers. Maximum intensity projections were generated in Zen Black, and unless otherwise stated, the images shown in this study represent these projections. Fluorescence intensity quantification was performed in FIJI/ImageJ (v1.53k). Regions of interest (ROI) corresponding to nuclei or expression domains were manually defined using standardized shapes and sizes across embryos, ensuring consistency within and across experimental conditions and developmental stages. Signal intensities were quantified from raw, single-channel images (not pseudocolor overlays). All quantification was performed blind to condition to avoid bias.

### Time-course quantification of HCR signal

To assess temporal changes in gene expression, HCR fluorescence was quantified from whole embryos across five embryos per time point (developmental stages 10-21) following a standardized thresholding workflow. Prior to measurement, each embryo was manually outlined using the polygon selection tool, and the background outside of the embryo was cleared to eliminate non-specific signals. Quantitation was then carried out using identical ROIs, positioned consistently across all embryos and timepoints. To isolate the relevant signal, fluorescence channels were manually thresholded to generate binary masks. Threshold levels were carefully adjusted to ensure signal detection without over- or under-thresholding. Mean gray values were recorded (arbitrary units, A.U.), and normalized fluorescence intensity values were calculated by dividing signal intensity (e.g., *Lmx1* or *Msx*) by DAPI intensity for each embryo. The per time point averages were plotted over time. The line chart was generated using GraphPad Prism; error bars represent the standard deviation across embryos at each time point. All images were processed using uniform parameters to ensure comparability across time points.

### Single-nucleus quantification of CTCF and statistical analysis

For high-resolution quantification of transcriptional output, corrected total cell fluorescence (CTCF) was calculated for individual nuclei using the formula:

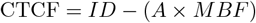

where *ID* is the integrated density, *A* is the area of the ROI, and *MBF* is the mean background fluorescence.

Scatter plots displaying the corrected total cell fluorescence (CTCF) of *Lmx1* and *Msx* over normalized y-coordinates were generated by measuring individual neural nuclei along the dorsal midline using a fixed-size circular ROI with an area of 32.09 px^2^. For each ROI, the mean gray value, integrated density, and coordinates were recorded for *n* = 3 embryos per time point (*n* = 4). T-tests were used to test the significance of CTCF between nuclei at the zippering point and nuclei either anterior or posterior to it. Statistical analysis (Welch’s t-test) was performed using the `Scipy.stats` function `ttest ind`. A ridgeline plot displaying posterior-to-anterior *Lmx1* transcription wave during neurulation was generated by recording the x- and y-coordinates of *Lmx1* -expressing nuclei and then normalizing the coordinates to the bounds of the embryo at respective time points. Graphs were generated using `matplotlib.pyplot`.

For dorsal midline cell number quantification, we counted the total number of *Lmx1::mnG* - or *Msx::mApp*- labeled nuclei per embryo (n = 20 embryos per condition, across n = 3 experiments) at 8 hours 27 minutes pf. A t-test was used to test significance between *Lmx1* or *Cdkn1b* overexpressing embryos and respective *LacZ* controls. Statistical analysis (student’s independent t-test) was performed using the `SciPy.stats` function `ttest ind`.

### scRNA-seq data analysis

Single-cell RNA sequencing (scRNA-seq) data from *Ciona intestinalis* type A (formerly *Ciona robusta*) initial tailbud I embryos (SRA accession numbers SRR9050993 and SRR9050994) were obtained from a previously published study (Cao et al., 2019). Raw reads were aligned to the latest *Ciona* genome assembly (HT) and the updated gene model (KY21) using 10x Genomics Cell Ranger (v2.0.1).

Post-alignment, single-cell data were analyzed using the Scanpy Python library (Wolf et al., 2018). Lowquality cells were filtered out by removing cells in which *>*20% of total counts were attributed to mitochondrial genes, *<*200 expressed genes, or a total number of unique molecular identifiers (UMIs) exceeding three standard deviations above the mean. Genes expressed in fewer than three cells were also excluded. Doublets were detected and removed using `Scrublet` (Wolock et al., 2019). Following filtering, 4,526 cells and 15,290 genes remained for downstream analysis.

Gene expression values were normalized to the median total count per cell using `scanpy.pp.normalize total`, followed by log-transformation with `scanpy.pp.log1p`. Highly variable genes were identified using `scanpy.pp.highly variable genes` with the Seurat v3 method (flavor = ‘seurat v3’, top 2,000 genes). These genes were scaled to unit variance and centered (values clipped at a maximum of 10) using `scanpy.pp.scale`.

Principal component analysis (PCA) was performed using the ARPACK solver. To correct for sample-specific batch effects, the top principal components were adjusted using the Harmony algorithm (Korsunsky et al., 2019), and the harmonized representation was stored in `X pca harmony`. A neighborhood graph was computed using `scanpy.pp.neighbors` on the Harmony-corrected components (n neighbors = 30, n pcs = 50).

Uniform Manifold Approximation and Projection (UMAP) was then applied for dimensionality reduction and visualization (McInnes et al., 2018). Clustering was performed using the Leiden algorithm (Traag et al., 2019) `scanpy.tl.leiden`, with various resolution parameters tested, including a high-resolution clustering (resolution = 10) to capture subpopulation heterogeneity.

## Author contributions

LAL, GÖ S and MSL conceptualized the research plan. GÖ S and LAL designed experiments, which GÖ S performed. GÖ S conducted image and data analysis and wrote the manuscript. LAL and MSL provided edits. MSL secured funding and supervised the work.

## Acknowledgments

We thank Dr. Nicholas Treen, Dr. Andrea Mariossi and Benjamin Larsen for helpful discussions, especially Dr. Nicholas Treen, who also shared reagents. We are also grateful for Dr. Harry McNamara and Dr. Emily Kolenbrander Ho for valuable feedback on the manuscript. The initial phases of this work were supported by a grant from the NINDS (NS076542) and completed with a grant from the Princeton Catalysis Initiative (PCI).

---

**Figure S1:**
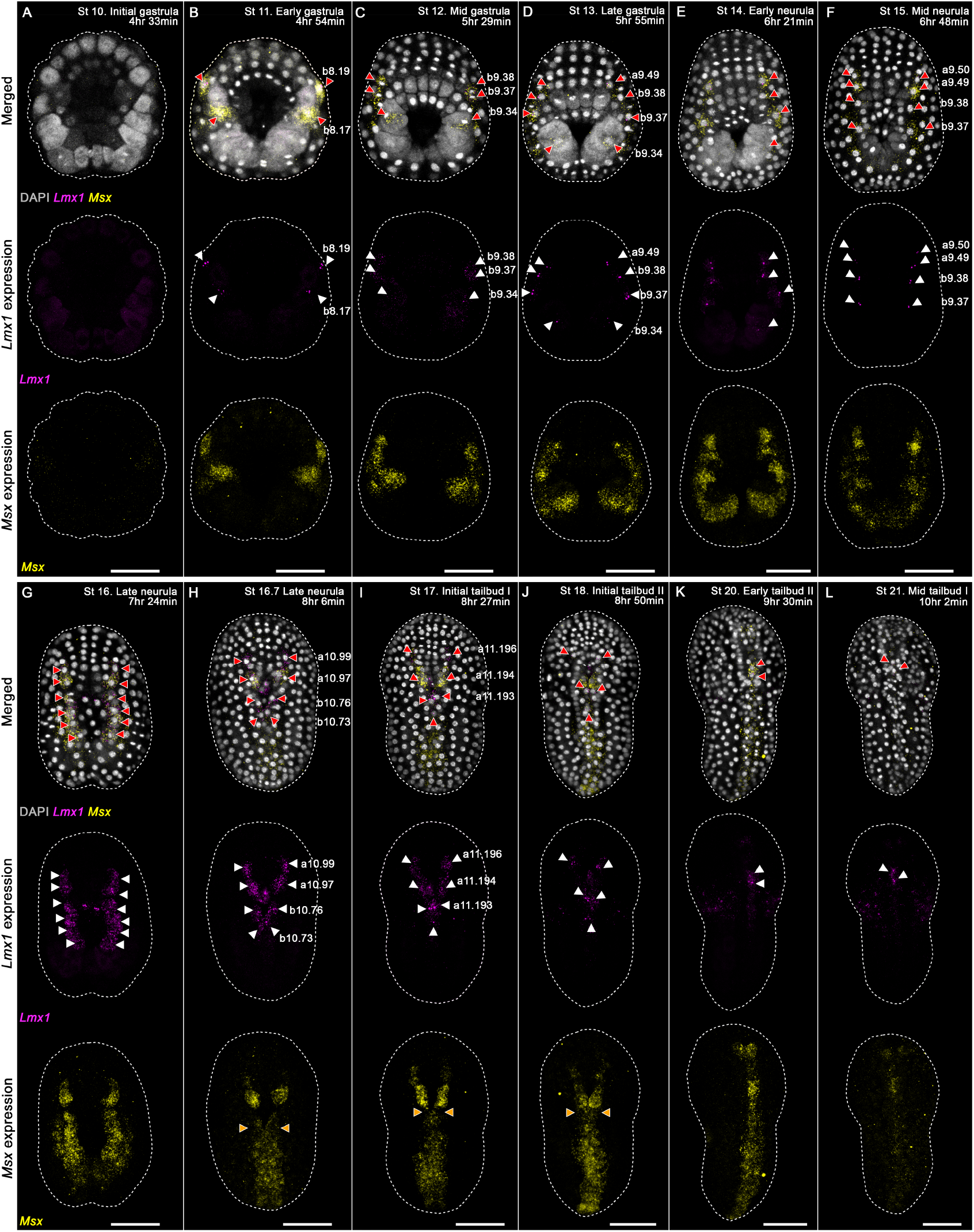
*Lmx1* and *Msx* expression patterns throughout gastrulation and neurulation. (A-L) Expression was examined throughout neurulation (∼4.5-10 hfp; samples collected every ∼20 minutes; 12 time points) via HCR *in situ* hybridization. The photographs are maximum-intensity projections of Z-projected image stacks overlaid in pseudocolor with HCR signals for *Lmx1* probe (magenta) and *Msx* probe (yellow). Nuclei were stained with DAPI (gray). Arrows (red/white) indicate *Lmx1* expression in descendants of neural plate border cells and neural plate cells, and (orange) *Msx* downregulation at zippering point. Brightness and contrast of images was adjusted linearly. Numbers of embryos examined n = 15 per time point over n = 5 experiments. Scale bars, 50 μm.

**Figure S2:**
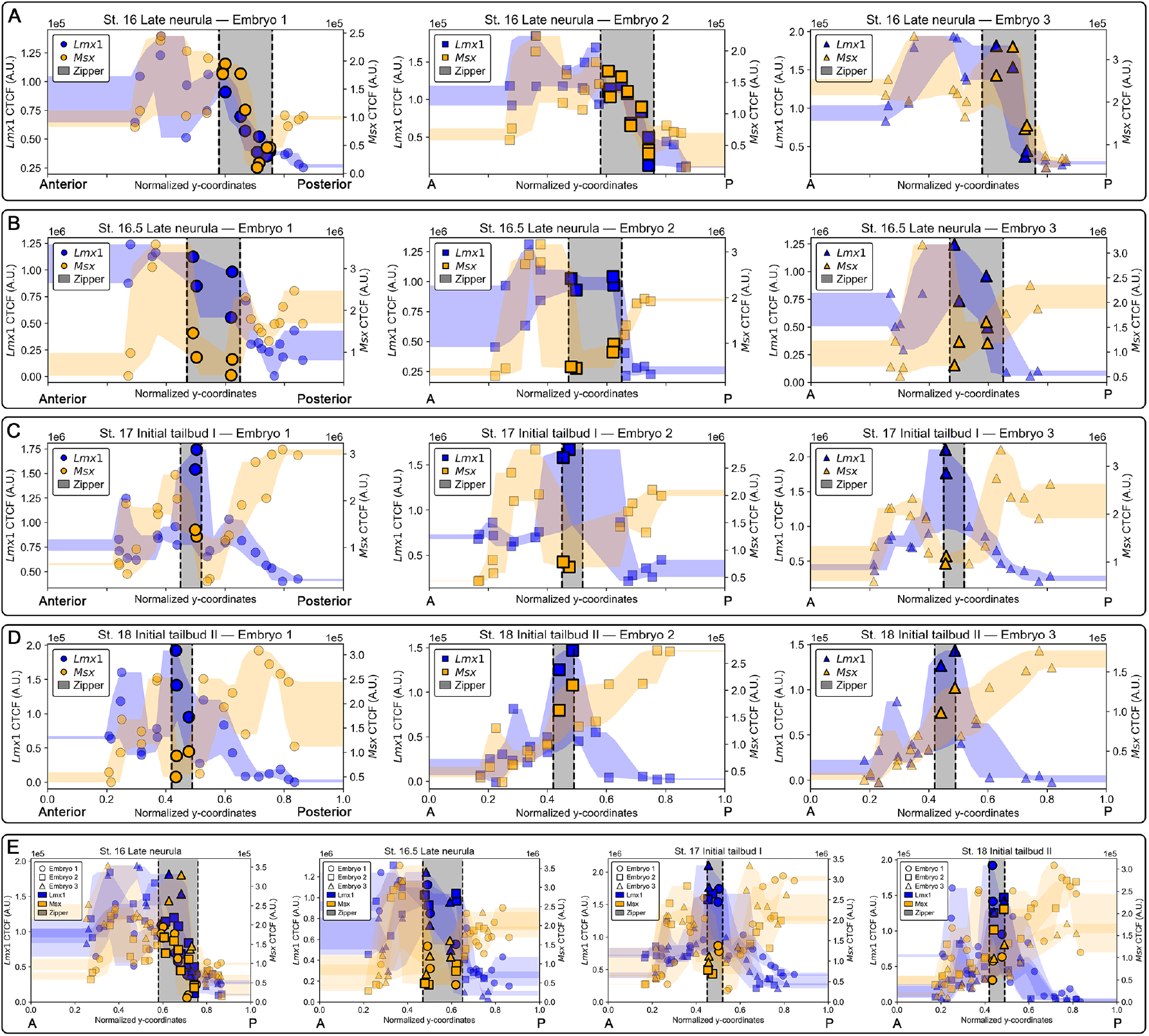
*Lmx1* expression increases while *Msx* is selectively downregulated at the zippering point during neural tube closure. (A–D) Corrected total cell fluorescence (CTCF) for *Lmx1* (dark blue) and *Msx* (orange) HCR *in situ* hybridization signals in dorsal midline cells across four developmental stages of NTC, plotted by each cell’s normalized y-coordinate (relative to the embryo height). Each point represents a single nucleus, and shaded regions denote the local range across neighboring points. Data are shown for n = 3 embryos per time point (n = 4), with marker shapes indicating individual embryos. During the Late Neurula stage (A), *Lmx1* and *Msx* are broadly co-expressed. As zippering initiates and progresses (B–C), *Lmx1* becomes locally enriched at the zippering point (gray region between dashed lines), while *Msx* becomes selectively downregulated in the same domain. (E) Aggregate plots from (A–D) illustrate consistency and embryo-to-embryo variability.

**Figure S3:**
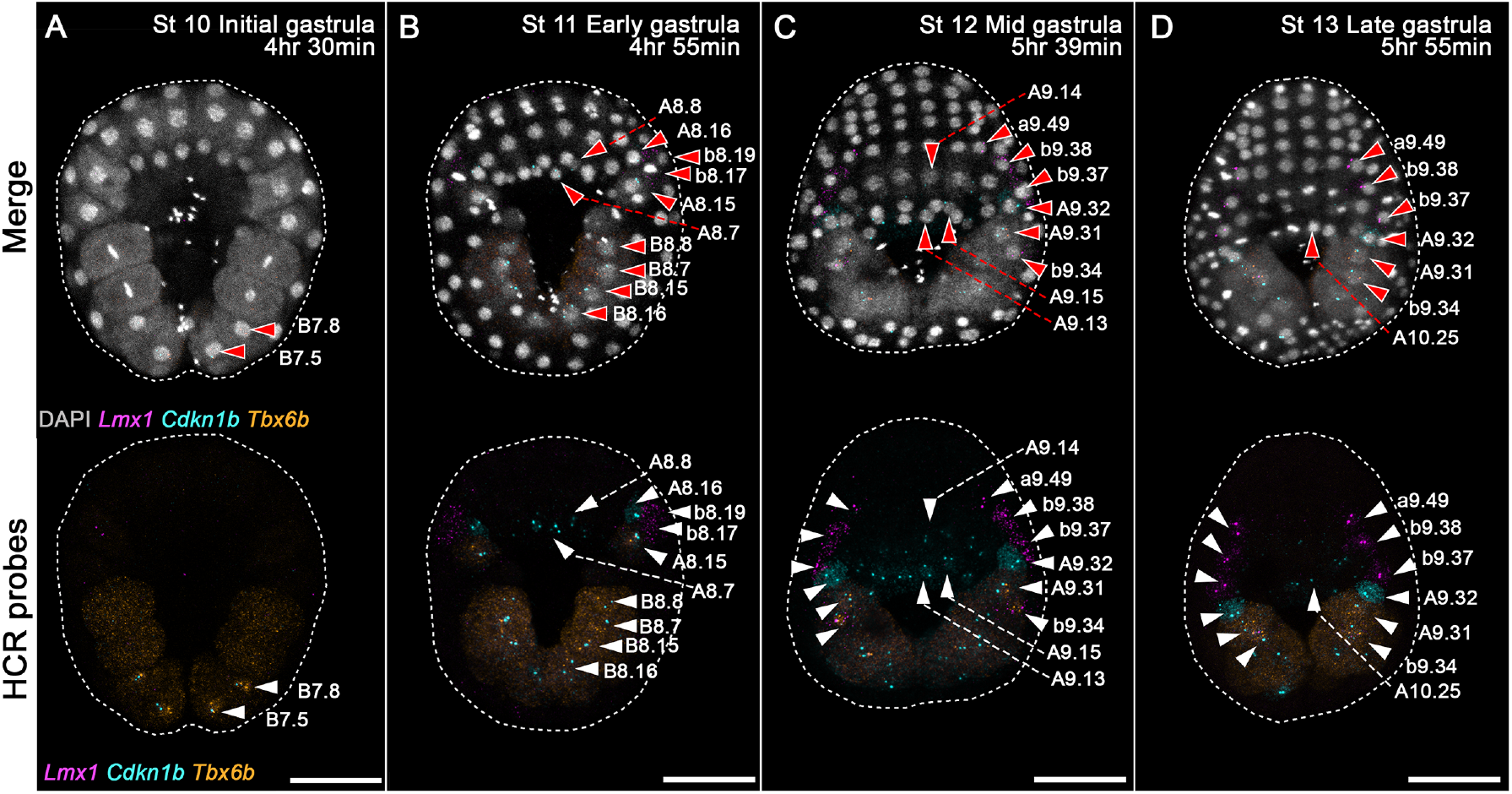
*Lmx1, Cdkn1b*, and *Tbx6-b* expression patterns during gastrulation. (A-D) Expression was examined throughout gastrulation (∼4.5-6 hfp; samples collected every ∼30 minutes; four time points) via HCR *in situ* hybridization. The photographs are maximum-intensity projections of Z-projected image stacks overlaid in pseudocolor with HCR signals for *Lmx1* probe (magenta), *Tbx6-b* probe (orange), and *Cdkn1b* probe (cyan). Nuclei were stained with DAPI (gray). Developmental stages are indicated in the photographs. Brightness and contrast of images was adjusted linearly. Numbers of embryos examined n = 15 per time point over n = 3 experiments. Scale bars, 50 μm.

**Figure S4:**
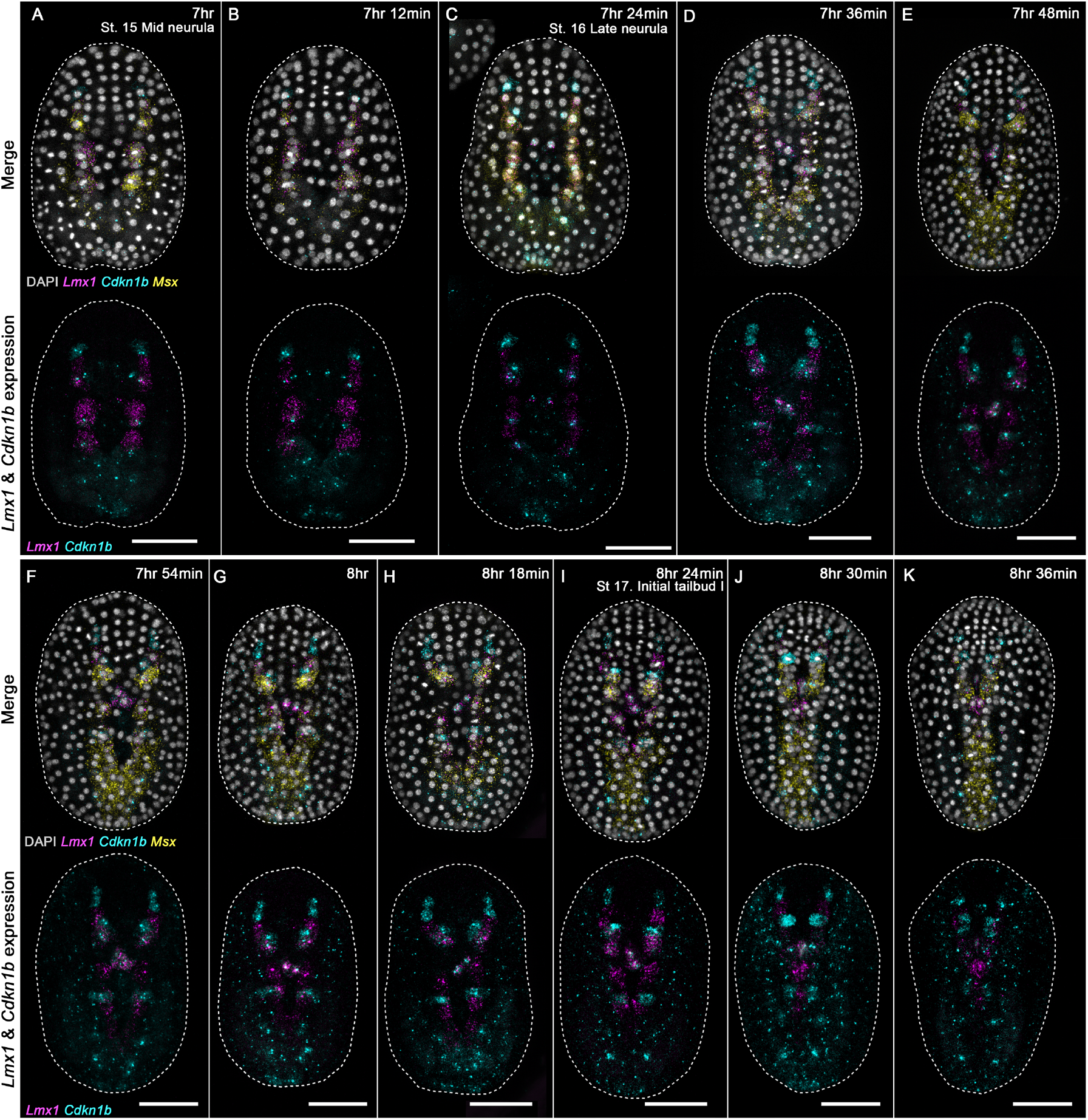
*Lmx1, Msx*, and *Cdkn1b* expression patterns during neurulation. (A-K) Expression was examined throughout neural tube closure (∼7-8.5 hfp; samples collected every ∼6 minutes; 13 time points) via HCR *in situ* hybridization. The photographs are maximum-intensity projections of Z-projected image stacks overlaid in pseudocolor with HCR signals for *Lmx1* probe (magenta), *Msx* probe (yellow), and *Cdkn1b* probe (cyan). Nuclei were stained with DAPI (gray). Developmental stages are indicated in the photographs. Brightness and contrast of images was adjusted linearly. Numbers of embryos examined n = 15 per time point over n = 3 experiments. Scale bars, 50 μm.

## References

Abitua, P. B., Wagner, E., Navarrete, I. A., and Levine, M. (2012). Identification of a rudimentary neural crest in a non-vertebrate chordate. Nature, 492(7427):104–107.

Angelini, T. E., Hannezo, E., Trepat, X., Marquez, M., Fredberg, J. J., and Weitz, D. A. (2011). Glass-like dynamics of collective cell migration. Proc. Natl. Acad. Sci. U. S. A., 108(12):4714–4719.

Avagliano, L., Massa, V., George, T. M., Qureshy, S., Bulfamante, G. P., and Finnell, R. H. (2019). Overview on neural tube defects: From development to physical characteristics. Birth Defects Res., 111(19):1455–1467.

Bocanegra-Moreno, L., Singh, A., Hannezo, E., Zagorski, M., and Kicheva, A. (2023). Cell cycle dynamics control fluidity of the developing mouse neuroepithelium. Nat. Phys., 19(7):1050–1058.

Cao, C., Lemaire, L. A., Wang, W., Yoon, P. H., Choi, Y. A., Parsons, L. R., Matese, J. C., Wang, W., Levine, M., and Chen, K. (2019). Comprehensive single-cell transcriptome lineages of a proto-vertebrate. Nature, 571(7765):349–354.

Chizhikov, V. V. and Millen, K. J. (2004). Control of roof plate formation by Lmx1a in the developing spinal cord. Development, 131(11):2693–2705.

Christiaen, L., Wagner, E., Shi, W., and Levine, M. (2009a). Electroporation of transgenic DNAs in the sea squirt ciona. Cold Spring Harb. Protoc., 2009(12):db.prot5345.

Christiaen, L., Wagner, E., Shi, W., and Levine, M. (2009b). Isolation of sea squirt (ciona) gametes, fertilization, dechorionation, and development. Cold Spring Harb. Protoc., 2009(12):db.prot5344.

Clark, E., Peel, A. D., and Akam, M. (2019). Arthropod segmentation. Development, 146(18):dev170480.

Copley, R. R., Buttin, J., Arguel, M.-J., Williaume, G., Lebrigand, K., Barbry, P., Hudson, C., and Yasuo, H. (2024). Early transcriptional similarities between two distinct neural lineages during ascidian embryogenesis. Dev. Biol., 514:1–11.

Dahlberg, C., Auger, H., Dupont, S., Sasakura, Y., Thorndyke, M., and Joly, J.-S. (2009). Refining the ciona intestinalis model of central nervous system regeneration. PLoS One, 4(2):e4458.

Fujiwara, D., Yoshimoto, H., Sone, H., Harashima, S., and Tamai, Y. (1998). Transcriptional co-regulation of saccharomyces cerevisiae alcohol acetyltransferase gene, ATF1 and delta-9 fatty acid desaturase gene, OLE1 by unsaturated fatty acids. Yeast, 14(8):711–721.

German, M. S., Wang, J., Chadwick, R. B., and Rutter, W. J. (1992). Synergistic activation of the insulin gene by a LIM-homeo domain protein and a basic helix-loop-helix protein: building a functional insulin minienhancer complex. Genes Dev., 6(11):2165–2176.

Greene, N. D. E. and Copp, A. J. (2014). Neural tube defects. Annu. Rev. Neurosci., 37:221–242.

Harris, M. J. and Juriloff, D. M. (2007). Mouse mutants with neural tube closure defects and their role in understanding human neural tube defects. Birth Defects Res. A Clin. Mol. Teratol., 79(3):187–210.

Harris, M. J. and Juriloff, D. M. (2010). An update to the list of mouse mutants with neural tube closure defects and advances toward a complete genetic perspective of neural tube closure. Birth Defects Res. A Clin. Mol. Teratol., 88(8):653–669.

Hashimoto, H. and Munro, E. (2019). Differential expression of a classic cadherin directs tissue-level contractile asymmetry during neural tube closure. Dev. Cell, 51(2):158–172.e4.

Hashimoto, H., Robin, F. B., Sherrard, K. M., and Munro, E. M. (2015). Sequential contraction and exchange of apical junctions drives zippering and neural tube closure in a simple chordate. Dev. Cell, 32(2):241–255.

Hotta, K., Mitsuhara, K., Takahashi, H., Inaba, K., Oka, K., Gojobori, T., and Ikeo, K. (2007). A web-based interactive developmental table for the ascidian ciona intestinalis, including 3D real-image embryo reconstructions: I. from fertilized egg to hatching larva. Dev. Dyn., 236(7):1790–1805.

Imai, K., Levine, M., Satoh, N., and Satou, Y. (2006). Regulatory blueprint for a chordate embryo. Science, 312:1183–1187.

Imai, K. S., Hino, K., Yagi, K., Satoh, N., and Satou, Y. (2004). Gene expression profiles of transcription factors and signaling molecules in the ascidian embryo: towards a comprehensive understanding of gene networks. Development, 131(16):4047–4058.

Irvine, K. D. and Wieschaus, E. (1994). Cell intercalation during drosophila germband extension and its regulation by pair-rule segmentation genes. Development, 120(4):827–841.

Ishida, T. and Satou, Y. (2024). Ascidian embryonic cells with properties of neural-crest cells and neuromesodermal progenitors of vertebrates. Nat. Ecol. Evol., 8(6):1154–1164.

José-Edwards, D. S., Kerner, P., Kugler, J. E., Deng, W., Jiang, D., and Di Gregorio, A. (2011). The identification of transcription factors expressed in the notochord of ciona intestinalis adds new potential players to the brachyury gene regulatory network. Dev. Dyn., 240(7):1793–1805.

Kancherla, V. (2023). Neural tube defects: a review of global prevalence, causes, and primary prevention. Childs. Nerv. Syst., 39(7):1703–1710.

Kim, K., Orvis, J., and Stolfi, A. (2022). Pax3/7 regulates neural tube closure and patterning in a non-vertebrate chordate. Front. Cell Dev. Biol., 10:999511.

Kim, S., Pochitaloff, M., Stooke-Vaughan, G. A., and Campàs, O. (2021). Embryonic tissues as active foams. Nat. Phys., 17(7):859–866.

Korsunsky, I., Millard, N., Fan, J., Slowikowski, K., Zhang, F., Wei, K., Baglaenko, Y., Brenner, M., Loh, P.-R., and Raychaudhuri, S. (2019). Fast, sensitive and accurate integration of single-cell data with harmony. Nat. Methods, 16(12):1289–1296.

Lehr, S., Brückner, D. B., Minchington, T. G., Greunz-Schindler, M., Merrin, J., Hannezo, E., and Kicheva, A. (2025). Self-organized pattern formation in the developing mouse neural tube by a temporal relay of BMP signaling. Dev. Cell, 60(4):567–580.e14.

McInnes, L., Healy, J., and Melville, J. (2018). UMAP: Uniform manifold approximation and projection for dimension reduction. arXiv [stat.ML].

Millonig, J. H., Millen, K. J., and Hatten, M. E. (2000). The mouse dreher gene Lmx1a controls formation of the roof plate in the vertebrate CNS. Nature, 403(6771):764–769.

Mishima, Y., Lindgren, A. G., Chizhikov, V. V., Johnson, R. L., and Millen, K. J. (2009). Overlapping function of Lmx1a and Lmx1b in anterior hindbrain roof plate formation and cerebellar growth. J. Neurosci., 29(36):11377–11384.

Moon, L. D. and Xiong, F. (2022). Mechanics of neural tube morphogenesis. Semin. Cell Dev. Biol., 130:56–69.

Negrón-Piñeiro, L. J., Wu, Y., Popsuj, S., José-Edwards, D. S., Stolfi, A., and Di Gregorio, A. (2024). Cis-regulatory interfaces reveal the molecular mechanisms underlying the notochord gene regulatory network of ciona. Nat. Commun., 15(1):3025.

Nikolopoulou, E., Galea, G. L., Rolo, A., Greene, N. D. E., and Copp, A. J. (2017). Neural tube closure: cellular, molecular and biomechanical mechanisms. Development, 144(4):552–566.

Nüsslein-Volhard, C. and Wieschaus, E. (1980). Mutations affecting segment number and polarity in drosophila. Nature, 287(5785):795–801.

Ogura, Y., Sakaue-Sawano, A., Nakagawa, M., Satoh, N., Miyawaki, A., and Sasakura, Y. (2011). Coordination of mitosis and morphogenesis: role of a prolonged G2 phase during chordate neurulation. Development, 138(3):577–587.

Ogura, Y. and Sasakura, Y. (2016). Developmental control of cell-cycle compensation provides a switch for patterned mitosis at the onset of chordate neurulation. Dev. Cell, 37(2):148–161.

Ramos, C. and Robert, B. (2005). msh/msx gene family in neural development. Trends Genet., 21(11):624–632.

Riddle, R. D., Ensini, M., Nelson, C., Tsuchida, T., Jesse ILl, T. M., and Tabin”, C. (1995). Induction of the LIM homeobox gene Lmx7 by WNT7a establishes dorsoventral pattern in the vertebrate limb. Cell, 83:631–640.

Roure, A. and Darras, S. (2016). Msxb is a core component of the genetic circuitry specifying the dorsal and ventral neurogenic midlines in the ascidian embryo. Dev. Biol., 409(1):277–287.

Sakamoto, R., Banerjee, D. S., Yadav, V., Chen, S., Gardel, M. L., Sykes, C., Banerjee, S., and Murrell, M. P. (2023). Membrane tension induces F-actin reorganization and flow in a biomimetic model cortex. Commun. Biol., 6(1):325.

Sasakura, Y., Mita, K., Ogura, Y., and Horie, T. (2012). Ascidians as excellent chordate models for studying the development of the nervous system during embryogenesis and metamorphosis. Dev. Growth Differ., 54(3):420–437.

Todorov, L. G., Oonuma, K., Kusakabe, T. G., Levine, M. S., and Lemaire, L. A. (2024). Neural crest lineage in the protovertebrate model ciona. Nature, 635(8040):912–916.

Traag, V. A., Waltman, L., and van Eck, N. J. (2019). From louvain to leiden: guaranteeing well-connected communities. Sci. Rep., 9(1):5233.

Treen, N., Chavarria, E., Weaver, C. J., Brangwynne, C. P., and Levine, M. (2023a). An FGF timer for zygotic genome activation. Genes Dev., 37(3-4):80–85.

Treen, N., Konishi, S., Nishida, H., Onuma, T. A., and Sasakura, Y. (2023b). Zic-r.b controls cell numbers in ciona embryos by activating CDKN1B. Dev. Biol., 498:26–34.

Wolf, F. A., Angerer, P., and Theis, F. J. (2018). SCANPY: large-scale single-cell gene expression data analysis. Genome Biol., 19(1):15.

Wolock, S. L., Lopez, R., and Klein, A. M. (2019). Scrublet: Computational identification of cell doublets in single-cell transcriptomic data. Cell Syst., 8(4):281–291.e9.

Yan, C. H., Levesque, M., Claxton, S., Johnson, R. L., and Ang, S.-L. (2011). Lmx1a and lmx1b function cooperatively to regulate proliferation, specification, and differentiation of midbrain dopaminergic progenitors. J. Neurosci., 31(35):12413–12425.

Zallen, J. and Wieschaus, E. (2004). Patterned gene expression directs bipolar planar polarity in drosophila. Dev. Cell, 6(3):343–355.

Zhao, J., Perkins, M. L., Norstad, M., and Garcia, H. G. (2023). A bistable autoregulatory module in the developing embryo commits cells to binary expression fates. Curr. Biol., 33(14):2851–2864.e11.

